# A simple, fast and cost-efficient protocol for ultra-sensitive ribosome profiling

**DOI:** 10.1101/2025.04.09.647970

**Authors:** Jiri Koubek, Katharina Jetzinger, Shiran Dror, Mikel Irastortza-Olaziregi, Dana Frank, Ilgin Kotan, Jaime Santos, Frank Tippmann, Pascal Lafrenz, Henrik Kaessmann, Orna Amster-Choder, Bernd Bukau, Günter Kramer

## Abstract

Ribosome profiling has become an essential tool for studying mRNA translation in cells with codon-level resolution. However, its widespread application remains hindered by the labour-intensive workflow, low efficiency and high costs associated with sequencing sample preparation. Here, we present a new cost-effective and ultra-sensitive library preparation method that significantly advances the applicability of ribosome profiling. By implementing bead-coupled enzymatic reactions and product purifications, our approach increases both yield and throughput while maintaining high reproducibility. Demonstrating the sensitivity of the protocol we prepared libraries from as little as 12 fmol of RNA, which expands the feasibility of ribosome profiling from minimal input samples, such as derived from small populations, stressed cells, or patient-derived specimens. Additionally, we validate the versatility of the protocol across multiple species and demonstrate its applicability for RNA-seq library preparation. Altogether, this protocol provides a highly accessible and efficient alternative to existing ribosome profiling workflows, facilitating research in previously challenging experimental contexts.

## Introduction

Ribosome profiling (RP) is a high-resolution sequencing-based method that provides a transcriptome-wide snapshot of active translation *in vivo*. By nuclease digestion of cellular mRNA followed by sequencing ribosome-protected mRNA fragments (footprints), RP enables precise mapping of translating ribosomes across the transcriptome. RP provides insights into gene expression levels, elongation dynamics, and translational regulation (1), making it a powerful tool to study translational pausing and frameshifting (2, 3), the effects of codon usage effects on protein synthesis (4), and uncovering novel open reading frames (5), among other processes. Furthermore, adaptations of RP, which focus on specific subsets of ribosomes, have been used to investigate a wide range of fundamental processes linked to protein biogenesis. These include co-translational complex formation (6–8), chaperone interactions (9), co-translational membrane targeting (10, 11), ribosome-associated quality control (12), and translation initiation (13). As a result, RP has become an essential tool for studying a broad spectrum of processes involved in protein biogenesis, with significant impacts on fields ranging from functional genomics to synthetic biology and disease modelling.

Despite its transformative potential, a major constraint limiting the widespread adoption of RP is the technically demanding nature of sequencing library preparation. This generally requires linker ligation and circularization reactions as well as multiple size selections of reaction products. Compared to standard RNA-seq, RP involves shorter fragment sizes (~25-40 nucleotides for ribosome-protected footprints). The size difference between the linkers and ligated products is several folds smaller, which makes the efficient purification of reaction products challenging (14). The classical RP library preparation protocol published by Ingolia and colleagues (1) relies on multiple PAGE purifications to isolate reaction products by size selection (Figure 1A). These purifications are not only time-consuming and prone to increasing the risk of accidents and cross-contamination but also significantly reduce material recovery (15), ultimately limiting the sensitivity of RP. Moreover, the use of circular ligase creates a financial bottleneck, taking up to 30 % of the library preparation costs (Table I).

**Figure 1:**
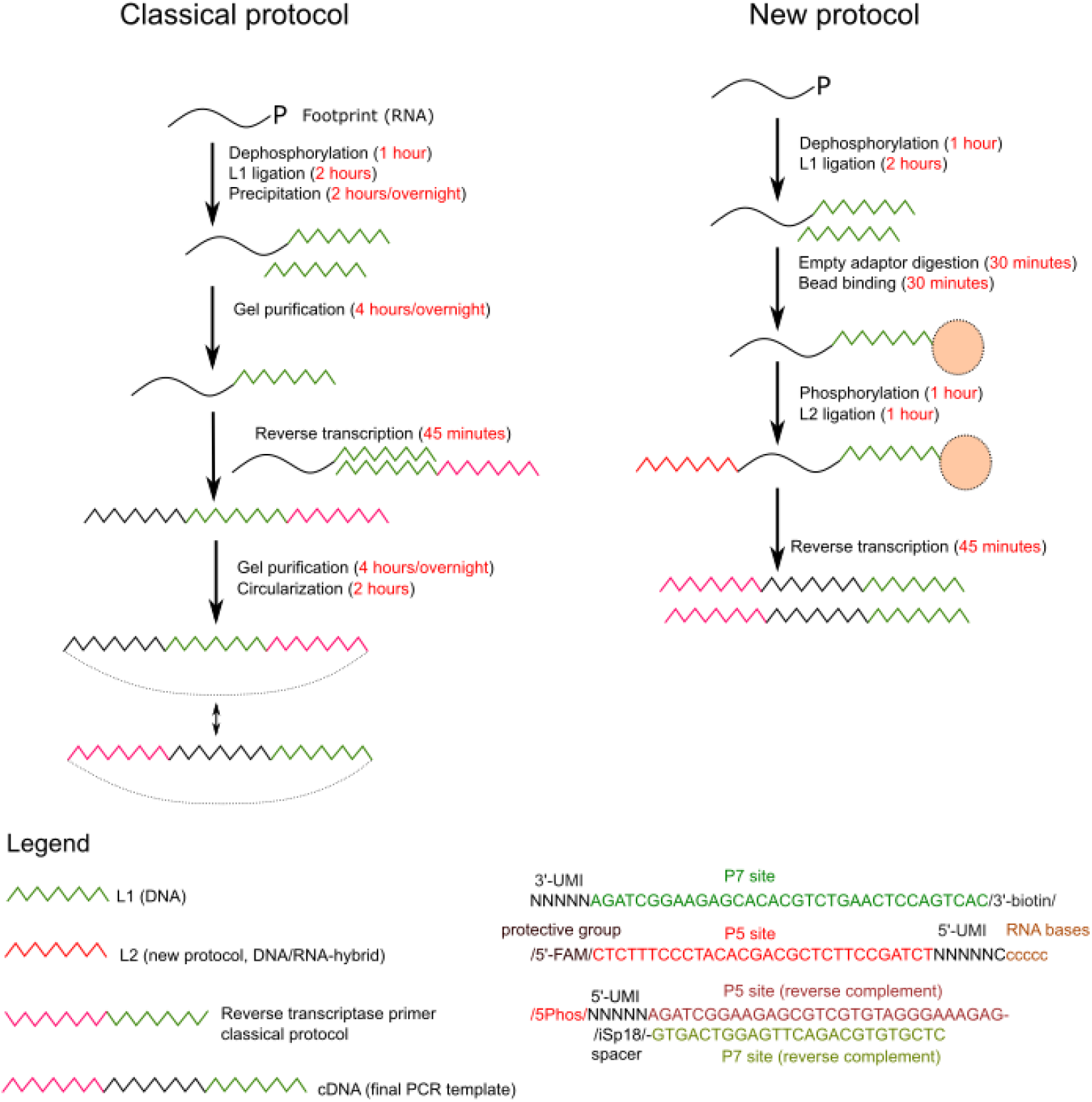
General overview of the new protocol

Over the years, the classical RP library preparation protocol has undergone several refinements (16, 17), along with the development of alternative methods that reduce workload while increasing sensitivity (15, 18, 19). Commercial library preparation kits have introduced alternative purification strategies to accelerate workflow, such as column or magnetic bead-based size selection. However, these come at the cost of lower purification yields and therefore lower sensitivity. Despite these developments, many labs continue to use the classical protocol, with minor variations, due to its well-established robustness, minimal sequence bias and high accuracy in footprint assignment.

Here, we present a highly streamlined and cost-effective RP library preparation protocol. Our new protocol introduces key optimizations, including the absence of circular ligation and the replacement of PAGE-based purification, enabling robust library preparation from substantially lower input material. This refined workflow not only enhances the sensitivity and scalability but also improves the accessibility of this technique to a broader range of laboratories. By reducing the costs and time while maintaining high data quality, this protocol expands the feasibility of RP for high throughput sequencing studies and challenging biological samples. This will contribute to diverse studies seeking deeper insights into translational control across diverse cellular and physiological contexts.

## Results

### General considerations and design

To streamline, simplify and reduce the cost of RP library preparation, we decided to focus on two of the major time and cost-consuming processes of the classical library preparation protocol; the recurrent use of PAGE-based purification and the circular ligation (Figure 1). The major changes which bypass the need for the two bottlenecks include the following: First, we introduced a biotinylated 3’ adaptor (L1), that allows biotin-streptavidin affinity purification of the ligation products, replacing the PAGE-based purification. Second, we optimised a 5’ adaptor (L2) ligation on beads, replacing the circularization step.

A brief overview of the individual steps of the new library generation is provided in Figure 1. Similar to the original protocol, ribosomal footprints are first dephosphorylated using polynucleotide kinase (PNK), followed by ligation of the 3’-biotinylated adaptor L1. The biotinylation serves two functions: (i) Ligation products can be coupled to streptavidin-coated magnetic beads, facilitating rapid and efficient purifications to replace PAGE-based purification. (ii) The biotin group prevents concatemer formation where multiple L1 adaptors are ligated to the same footprint. Removal of unreacted L1 is facilitated by incubation with 5’ deadenylase and DNA exonuclease (17). Next, the 5’-end of bead-captured products are phosphorylated with PNK. Separating the 5’-end phosphorylation reaction from the preceding 3’-end dephosphorylation reaction greatly improves the L1 ligation (Figure S1). Phosphorylated products are ligated with linker L2. L2 is a 5’-fluoresceine (5’-FAM)-labeled DNA-RNA hybrid (Figure 1) and ligation to the 5’-end of the footprint is performed with T4 RNA ligase I (20), which eliminates the need for circularization and greatly reduces the costs of library preparation. Using a DNA-RNA hybrid protects the 5’-end from degradation by contaminating RNA 5’-3’ exonucleases and lowers the chance of random nucleophilic attack of the 2’-OH group which is innate to all RNA molecules (21). Subsequently, the products are reverse-transcribed and library preparation is completed by a small number of PCR cycles (Figure 1) to amplify the fragments and incorporate Illumina sequencing-specific P5 and P7 sites.

### Protocol validation

To validate our new protocol, we conducted side-by-side comparisons with the classical method using sucrose cushion-purified ribosomes isolated from the same nuclease-treated *E. coli* lysate. Gel-extracted footprints (5 μg of total RNA) were split equally into two parts and libraries were prepared with either protocol. Both libraries were sequenced to similar depths; reads were aligned to the *E. coli* genome and the reproducibility was determined by comparing the footprint distributions along coding sequences.

The results confirm that our new protocol fully replicates the data quality of the classical approach (Figure 2). We observed comparable yields of the expected full-length fragments between the protocols (Figure 2A) and found strong correlation in total read count per gene (Figure 2B). Notably, the footprint distributions along coding sequences – which reflect local translation dynamics – was highly consistent between the two methods, as exemplified for the gene *rpsA* (Figure 2C). Furthermore, the individual gene profile reproducibility is detectable for genes with high coverage genome-wide (Figure 2D). Together these data demonstrate that the new protocol can fully substitute the high-quality output of the classical protocol for only ~30 % of both costs and time (Table 1).

**Table 1:** Comparison of costs and time required for each library preparation protocol, only incubation times are listed.

|  | Classical protocol |  | New protocol |  |
| --- | --- | --- | --- | --- |
|  | time | Costs* | time | Costs* |
| Footprint extraction† | 4 hours/overnight | 13 | 4 hours/overnight | 13 |
| Dephosphorylation | 1 hour | 1.8 | 1 hour | 0.7 |
| L1 ligation | 2 hours | 5.3 | 2 hours | 2.1 |
| Precipitation | 2 hours/overnight | 0.4 |  |  |
| Gel extraction† | 4 hours/overnight | 7.7 |  |  |
| L1 depletion |  |  | 30 minutes | 3.9 |
| Bead binding |  |  | 20 minutes | 0.4 |
| Phosphorylation |  |  | 1 hour | 0.7 |
| L2 ligation |  |  | 1 hour | 0.8 |
| Reverse transcription | 45 minutes | 8.3 | 45 minutes | 8.3 |
| Gel extraction† | 4 hours/overnight | 7.7 |  |  |
| Circularization | 2 hours | 41.2 |  |  |
| Final PCR‡ | 1 hour | 1.3 | 1 hour | 1.3 |
| **Total** | **5 days** | **86.7** | **2 days** | **31.2** |

\*costs (in euros) were calculated using reagents retail values available in Germany in February 2025
†Includes costs of precast gel, gel breaker tubes, SpinX columns and glycoblu
‡PCR purification may include additional PAGE-purification and precipitation.

**Figure 2:**
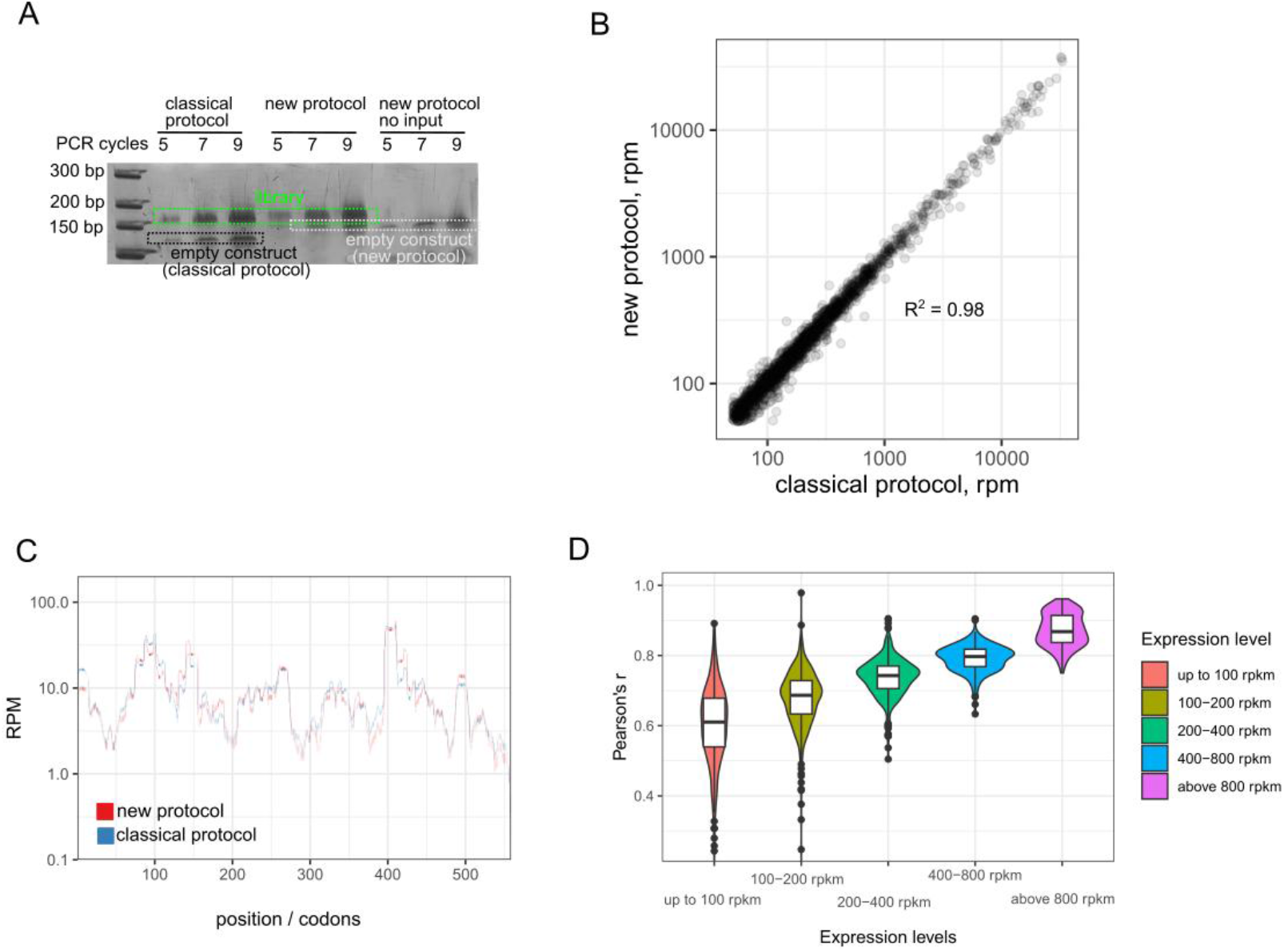
New protocol reproduces results from classical protocol in *E. coli* RP. A) PCR amplification gel, B) Comparison of detected gene expression levels between the classical protocol and the new protocol. Linear reproducibility R2 is calculated from log-transformed rpm values. C) Ribosome profile of *rpsA* smoothed over 15 codons. D) Overall peak height reproducibility between the classical and new protocols genome-wide. Genes are sorted according to their expression levels.

### The new library protocol has superior sensitivity

To determine the sensitivity of the protocol, we prepared sequencing libraries from serial dilutions of 28 nt control RNA oligonucleotide, ranging from 12 fmol to 1000 fmol of input material. We then evaluated the number of PCR cycles required for generating DNA fragments containing the input material. While in the classical protocol, about 1000 fmol of input material was needed to generate a library after 5 PCR cycles, the new library only needed 37 fmol of input material to get the same result (Figure 3A). To quantify the higher sensitivity, we calculated the RNA conversion rate of both protocols, i.e. the total yield of amplifiable cDNA from input RNA. Using control RNA oligonucleotide, the classical protocol yielded a conversion rate of 2 %, as compared to a conversion rate of about 30 % using the new protocol. Most notably, only 7 PCR cycles were needed to amplify libraries from 12 fmol of input material. This amount of input material can be recovered from about 30 ng of *E. coli* ribosomes (22).

**Figure 3:**
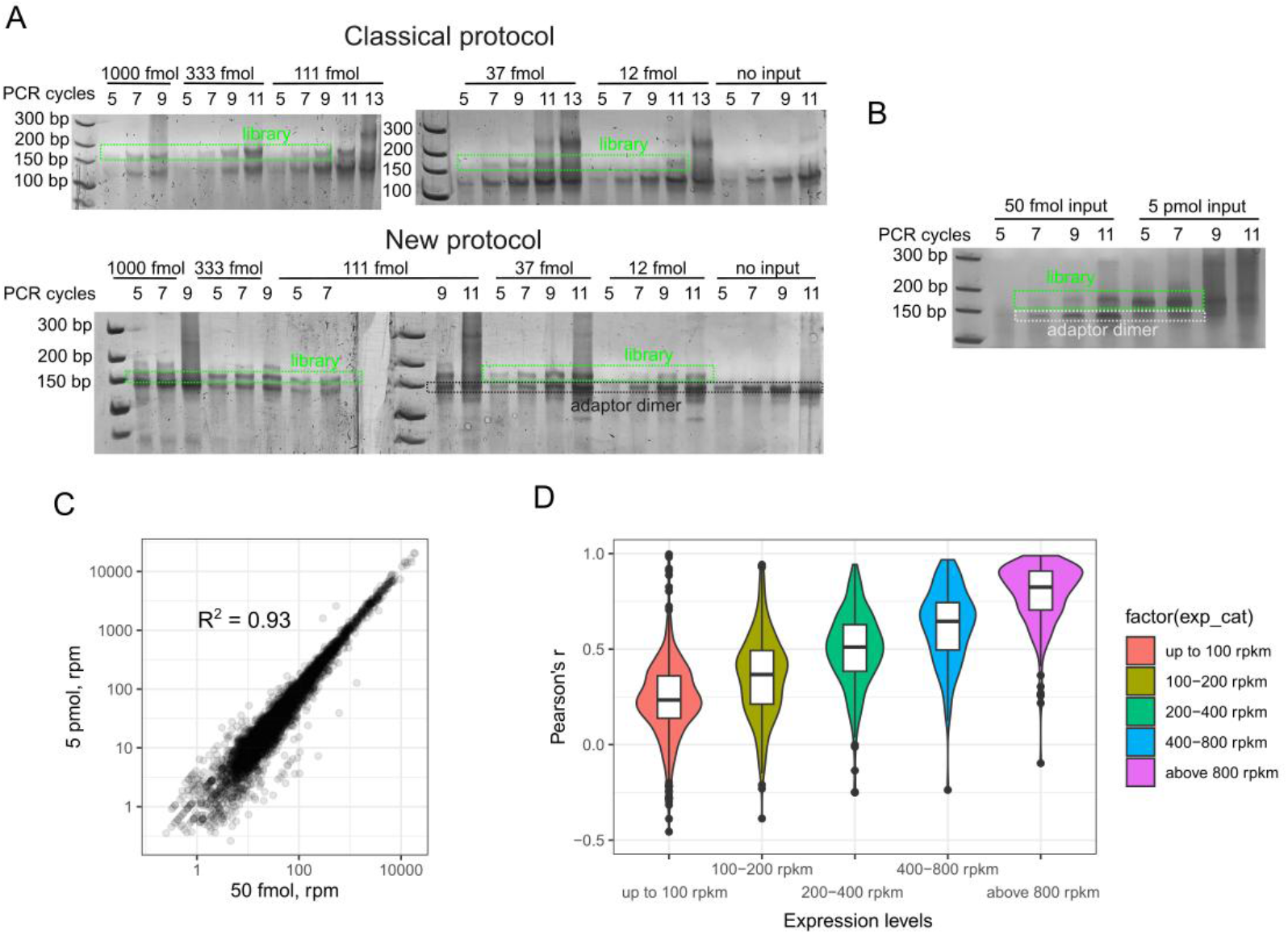
The new protocols offers up to 12 fmol input material sensitivity. A) Comparison of PCR amplification step at different amount of inputs using a defined control oligo. cDNA was prepared with either the classical protocol or the new protocol. B) PCR amplification step of S. cerevisiae total translatome libraries using 50 fmol or 5 pmol input. C) Gene expression reproducibility of different inputs of yeast total translatome. Linear reproducibility R2 is calculated from log-transformed rpm values. D) Overall peak height reproducibility between the 5 pmol and 50 fmol input genome-wide, peaks were calculated as a sliding window over 5 codons. Genes are sorted according to their expression levels.

Next, we validated the protocol sensitivity by generating total translatome libraries from *S. cerevisiae* footprints, using 5 pmol or 50 fmol of gel-extracted RNA fragments (Figure 3B), which corresponds to a yield from roughly 10 μg or 100 ng ribosomes, respectively (22). Confirming the protocol faithfully captures the translatome independently of the input levels, the bioinformatic analyses of the sequenced libraries revealed highly correlated gene expression levels (Figure 3C) and footprint distributions along gene profiles were well-correlated among highly covered genes (Figure 3D). We conclude that the new protocol is at least 10-fold more sensitive than the classical protocol without compromising library complexity, quality or accuracy. This enhanced sensitivity expands the applicability of RP to low-yield samples, such as rare cell populations and challenging experimental conditions.

### Mitigating empty construct formation and rRNA content in low input samples

As observed during testing the sensitivity of the new protocol, preparing libraries from very low amounts of input material sometimes generates high proportions of L1-L2 dimer product (empty construct). L1-L2 dimers can affect sequencing yields if the final PAGE purification is insufficient. While this issue can be mitigated by reducing the amount of L1 in the first ligation reaction, we also aimed to develop a more straightforward strategy to improve the quality of low input libraries. For this purpose, we redesigned the sequence of L1 to create a uniform site when L1 and L2 are directly ligated. This was achieved by relocating the 5 random unique molecular identifier (UMI) nucleotides from L1 to L2 (Figure 4A). The uniform site formed by the undesired ligation can be bound by an added antisense oligo to selectively block reverse transcription, using locked nucleic acids (LNA) for higher specificity (23). In addition, the overhang of LNA on L1 without footprint will block L2 ligation as it will imitate nick joining on RNA duplexes which is inefficient for T4 RNA ligase 1 (24) (Figure 4B). Overall, we observed that the presence of the LNA in the L2 ligation and reverse transcription led to a 6-fold reduction of empty construct in the final library (Figure 4C and Table 2). Although this modification slightly complicates the protocol and moderately raises the costs, it is highly recommended for high-throughput applications where the amount of L1 cannot be adjusted to the amount of the input material.

**Table 2:** Reduction of adaptor-dimers in sequenced libraries.

|  | No LNA | ligation | RT | Ligation + RT |
| --- | --- | --- | --- | --- |
| Dimers | 61276 | 28150 | 18564 | 15750 |
| rRNA | 142594 | 186630 | 180035 | 184463 |
| Genome-aligned | 145358 | 210791 | 200350 | 214362 |
| % dimers to genome-aligned | 42 | 13 | 9 | 7 |

\*values normalized to 1M sequenced reads

**Figure 4:**
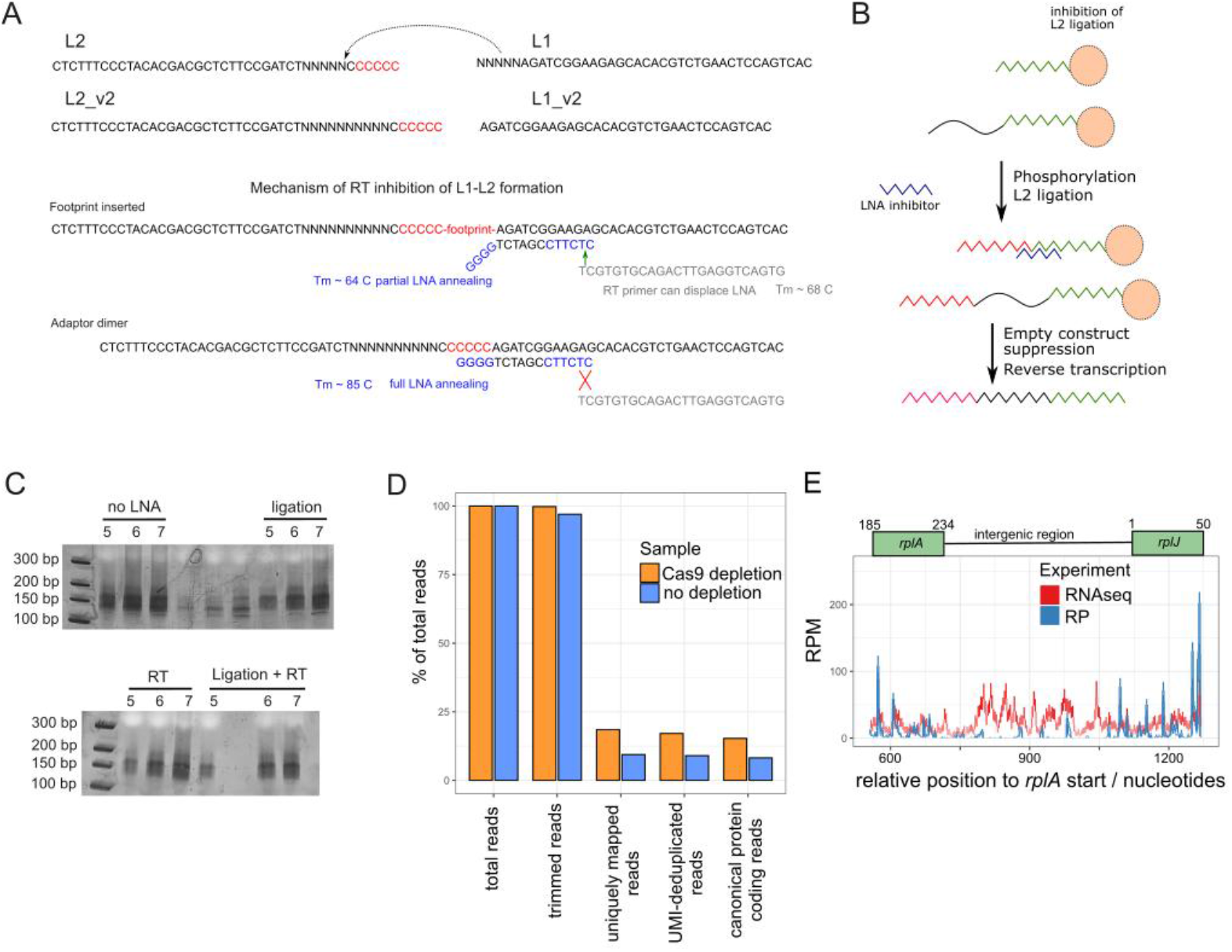
Increasing the utility of the new protocol a) Scheme of redesigned linkers for LNA-based empty construct depletion, b) Adaptation of the protocol, c) Gel of the PCR amplification step using LNA at various steps of library preparation, d) Implementing Cas9-based rRNA depletion leads to 2-fold enrichment of reads in protein-coding regions, e) RNAseq prepared by the new protocol can cover intergenic regions of the *rplKAJL* operon.

The presence of contaminating rRNA fragments is one of the major limitations in RP, often comprising up to 90 % of sequencing reads. Traditional depletion strategies rely on biotinylated antisense oligonucleotides targeting rRNA, either during the footprint isolation stage or before cDNA preparation (25). However, for low-input samples, performing rRNA depletion at the footprint stage is not feasible, as it significantly reduces the already limited input material, rendering library preparation impossible. We therefore evaluated the effectiveness of recently described Cas9-based rRNA depletion strategies (26), which removes PCR fragments derived from rRNA, for RP of mouse brain samples. In this case, the input is no longer limiting, as the depletion is performed after the PCR amplification step. Our results showed that the proportion of reads mapping to protein-coding regions nearly doubled (Figure 4D), suggesting that Cas9-based depletion could be a valuable addition to our protocol, further enhancing sequencing efficiency.

### Adaptability to other sequencing applications

Finally, we explored whether our protocol can also be applied to other sequencing applications, such as RNA-seq. By applying minimal modifications to obtaining the starting RNA fragments, we successfully prepared standard RNA-seq libraries (Figure 4E). Unlike in RP, RNA-seq provides a coverage for the entire transcripts which in bacteria include intergenic regions in operons. Comparing data of RNA-seq to RP, we observed a well-covered intergenic region between the *E. coli* genes *rplA* and *rplJ* which are transcribed in the same operon, demonstrating its broader utility beyond RP.

## Discussion

RP has emerged as a powerful method for studying translation with nucleotide resolution, but its application has been limited by the need for substantial input material and labor-intensive library preparation protocols. Over the past decade, numerous refinements have been introduced to improve the efficiency (15, 17–19), sensitivity, and accessibility of RP, yet significant challenges remain in adapting the method for low-input applications. Our study presents a novel library preparation protocol that overcomes several of these limitations. It enables RP from as little as 12 fmol of input material with a footprint conversion rate of 30 % (from input material to sequencing library), increasing the sensitivity by at least one order of magnitude. By integrating on-bead reactions, we provide a platform for high throughput applications, minimizing handling while maintaining data reproducibility.

Several studies have sought to enhance RP for low-input applications. To our knowledge, the highest sensitivity is provided by the recent protocol developed by Meindl et al. (15). It introduced optimizations that reduce sample handling time while maintaining detection accuracy, as complex sequencing libraries could be prepared from as little as 100 fmols. However, their method still relies on circular ligation, which remains a significant bottleneck due to its high costs. Similarly, Ferguson et al. (18) proposed a single-tube sequencing workflow using a two-template relay strategy (27) to enhance ligation efficiency. While effective, this approach depends on specialized enzymatic reactions that may not be broadly accessible, making it less universally adaptable. In contrast, our protocol uses common T4 RNA ligases to avoid fragment circularization and applies a streptavidin bead-based purification to replace gel-based purifications. In consequence the protocol is only limited by the efficiency of the enzymatic reactions. This results in exceptionally high footprint conversion rates (30 % compared to 2 % in the classical protocol) while also significantly cutting the costs of library preparation without compromising data complexity and quality.

By significantly reducing input material requirements and streamlining library preparation, our method expands the applicability of RP to biological contexts where sample availability is limited. This includes the analysis of rare cell types, which is out of reach for less sensitive traditional approaches, as well as samples from early developmental stages, which have so far not been studied with conventional methods. Additionally, our protocol enhances the feasibility of studying translation under stress conditions, such as nutrient deprivation or oxidative stress, where low translation rates (28, 29) often limit footprint recovery. Moreover, as interest in RP grows in clinical and translational research, our approach provides a practical solution for analysing changes of mRNA translation in patient-derived samples, facilitating its potential applications in personalized medicine. While progress has been made on single-cell RP (20, 30), current methods require highly specialized equipment, reagents and protocols. The advances described in this study may serve as a foundation for improving these methods, allowing for more scalable and cost-effective single-cell applications. Lastly, with only very few modifications, we could successfully prepare RNAseq libraries, suggesting a broader utility in sequencing. We surmise our protocol can be used for the generation of any other short RNA-seq library, in particular those where sample size is a concern, such as CLIP-seq (31), NET-seq (32), or miRNA seq.

## Acknowledgement

This work was supported by the German-Israeli Foundation for Scientific Research and Development (I-1564-417.13/2023) to G.K. and O.A-C. and grant KR 3593/4-1 of the Deutsche Forschungsgemeinschaft to G.K. and B.B.. B.B. acknowledges a research grant of the European Union (ERC-SyG-101072047-CoTransComplex). Views and opinions expressed are however those of the authors only and do not necessarily reflect those of the European Union or the European Research Council. Neither the European Union nor the granting authority can be held responsible for them. We furthermore thank the Klaus Tschira Foundation for their generous support in the acquisition of the sequencing device. JK was supported by European Research Council (Horizon 2020 Individual Fellowship, CoCoAssembly 895164). D.F., P.L., and H.K. were supported by a NOMIS Foundation Researcher Grant (no. 2024-A-0035, to H.K.). MIO’s stay in Heidelberg was supported by the GIF (I-1564-417.13/2023).

## Materials and methods

### Used media and strains

We used wild type *E. coli* MG1655 strain and *S. cerevisiae* BY4741, unless specified otherwise. *E. coli* were grown on LB media at 37 °C unless specified otherwise. *S. cerevisiae* cells were grown in YPD media (1 % yeast extract, 2 % peptone, 2 % glucose) at 30 °C.

### Cell harvest and lysis

*E. coli* cells were grown to OD600 0.3-0.5 with shaking at 37 °C in 200 ml LB media. Cells were harvested by rapid filtration by passing through 0.2 μm nitrocellulose membrane as described earlier unless specified otherwise. Frozen cell pellets were mixed with 500 μl *E. coli* lysis buffer (50 mM Tris pH 7.0, 150 mM KCl, 10 mM MgCl_2_, 5 mM CaCl_2_ 0.1 % NP-40, 0.02 % Triton-X, 1x EDTA free protease inhibitor tablet (Roche), 0.02 U/μl DNaseI (Roche) and 1 mM chloramphenicol) via mixer milling (2 min, 30 Hz, Retsch).

For *S. cerevisiae* samples, 200 ml YPD media was inoculated with overnight cultures at an OD of 0.03 and grown to early log phase (OD600 0.4-0.5) with shaking at 30 °C. Cells were rapidly filtrated through nitrocellulose membranes with a pore size of 0.4 μM, scraped with a metal spatula and frozen in liquid nitrogen. Frozen cell pellets were lysed together with frozen droplets of yeast lysis buffer (20 mM Tris pH 8.0, 140 mM KCl, 10 mM MgCl_2_, 0.1 % NP-40, 0.1 mg/mL Cycloheximide, EDTA free protease inhibitor tablet (Roche), 0.02 U/μl DNaseI (Roche) and 40 μg/mL bestatin) via mixer milling (2 min, 30 Hz, Retsch).

### Ribosome isolation

For *E. coli*, lysates were thawed by addition of 500 μl of *E. coli* lysis buffer. RNA concentration was determined by Nanodrop and RNA was digested with Micrococcal nuclease (33) using 150 U per 40 μg of RNA at 25 C for 10 minutes. The reaction was stopped by adding 6 mM EGTA (final concentration) and the lysates were clarified by centrifugation at 20 000 x g for 3 minutes. Clarified lysates (200 μl) were overlaid on 600 μl sucrose cushion (30 % sucrose, 50 mM HEPES, 150 mM KCl, 10 mM MgCl2) and centrifuged in S120-AT2 rotor (Thermo Scientific) for 1 hour at 100 000 rpm. Supernatant was discarded, the pellet was resuspended in 100 μl of lysis buffer and subsequently extracted using Trizol reagent (Zymo research).

For *S. cerevisiae*, lysate was thawed for 2 min in a 30°C water bath. RNA concentration was measured using NanoDrop and RNA digestion was carried out for 30 minutes at 4°C, with 10 U RNase I per 40 μg of RNA. Digestion was stopped by addition of 10 μL Superase-In (Ambion). Lysates were clarified by centrifugation at 500 g for 5 min at 4 °C. The clarified lysate was layered on a sucrose cushion (25 % w/v sucrose, 20 mM Tris HCl pH 8.0, 140 mM KCl, 10 mM MgCl_2_, 0.1 mg/mL Cycloheximide, 1X Roche EDTA free complete PI tablets, 40 μg/mL Bestatin) and centrifuged for 90 min at 75,000 rpm at 4 °C in an S120-AT2 rotor (Thermo Scientific). The ribosomal pellet was resuspended in 700 μL of lysis buffer and RNA was extracted via Phenol-chloroform extraction (22).

### Footprint isolation

Extracted RNA from purified ribosomes was separated on a 15% denaturing urea polyacrylamide gel (Thermo Fisher Scientific) and stained with SYBR-Gold (Thermo Fisher Scientific). Gel slices with footprints with sizes 20–40 nt were excised and extracted using modified crush and soak method. Briefly, gel pieces were fragmented using gel breaker tubes. RNA was extracted in 0.5 mL of 10 mM Tris-HCl (pH 7.0) for 15 min at 75 °C with agitation. Gel debris was removed by centrifugation in Spin-X filter tubes (Corning) for 2 min at 15 000 x g. RNA was precipitated overnight at −70°C in the presence of 1 volume μL isopropanol, 0.1 volume 3 M sodium acetate (pH 5.5), and 2 μL GlycoBlue (Thermo Fisher Scientific). RNA was pelleted by centrifugation for 40 min at maximum speed, washed with 80% ethanol.

### Sequencing library preparation

For libraries prepared by the classical protocol, the samples were handled as described (34).

Isolated footprints were dephosphorylated by resuspending the washed RNA pellet with 4 μl PNK reaction solution (1 x PNK buffer, 3.2 U PNK, 16 U Murine RNAse inhibitor) and incubated for 1 hour at 37 C. PNK was inactivated by incubating at 75 C for 10 minutes. 4 μl of L1 ligation mixture (1x RNA Ligase Buffer, 1 μM pre-adenylated L1, 80 U T4 RNA Ligase 2 truncated, and 30 % PEG-8000) was added into the reaction. The resulting mixture was incubated at 22 C for 2 hours after which the unligated L1 was digested by adding 25 U deadenylase and 45 U of RecJf and incubated at 37 C for 30 minutes. The reaction was quenched by incubating at 75 C for 10 minutes.

Each reaction was then incubated for 10 minutes with 2 μl of magnetic streptavidin Dynabeads preequilibrated in 50 μl of BW buffer (10 mM Tris, pH = 7.0, 500 mM NaCl, 1 mM EDTA, 0.05 % Tween-20). After magnetic separation, supernatant was discarded and beads were washed two times with 200 μl of BW buffer and two times with 200 μl mild wash buffer (10 mM Tris, pH = 7.0, 0.05 % Tween-20) at 50 C. Washed beads were resuspended in 4 μl of phosphorylation reaction (1x PNK buffer, 3.2 U of PNK, 1 mM ATP, and 0.6x mild wash buffer) and incubated at 37 C for 1 hour. The reaction was then mixed with 4 μl of L2 ligation mixture (1x RNA Ligase Buffer, 1 μM L2, 1 mM ATP, 12 U RNA Ligase 1, and 30 % PEG-8000) and incubated at 37 C for 1 hour.

After magnetic separation, supernatant was discarded, the beads were washed once with 50 μl of mild wash buffer and resuspended in 8 μl of mild wash buffer. RT primer was annealed by incubating the beads at 65 C with 3 uL of RT1 mixture (6.7 uM RT primer, 3.3 mM each dNTP). After 5 minutes, the beads were placed on ice and mixed with 5 μl of RT2 mixture (3 μl of First strand buffer, 1 μl 100 mM DTT, 1 μl SuperScript III) and incubated at 50 C for 30 minutes. After incubation, the beads were placed at room temperature and magnetically separated. The supernatant was removed, the beads were washed once with 50 μl of mild wash buffer and subsequently resuspended in 30 μl of mild wash buffer.

### Final library preparation and deep sequencing

Samples containing cDNA (from classical and new protocol) were amplified by PCR. Each PCR amplification reaction contained the following: 4 μl HF Phusion Plus buffer (Thermo), 3 μl RT product containing beads, 0.4 μl fwd Primer (20 μM), 0.4 μl barcoded reverse Primer (20 μM), 0.4 μl 10 mM dNTPs, 11.6 μl ddH2O, and 0.2 μl Phusion Plus Polymerase (Thermo). The product was amplified for 5-11 cycles and resolved on 6 % PAGE-TBE gel run with 0.5 μg of Ultra Low Range DNA ladder. The product was quantified using ImageJ ((35)) using 150 bp and 200 bp marker bands as standards (each containing 30 ng of DNA). Footprints-containing PCR products (~160-180 bp) were excised and extracted by modified crush and soak method as described in the section on footprint isolation.

Isopropanol precipitated samples were resuspended in 12 μl 10 mM Tris (pH = 7.0), quantified using Qubit HS DNA assay, and fragment length was analysed on Bioanalyzer using dsDNA High sensitivity chip. Samples were pooled and their concentration was adjusted to 2-4 nM (final multiplex) and sequenced in house on Illumina NextSeq 550 using the NextSeq 500 High Output v2.5 Kit (75 cycles) with 6 bp i7 barcodes.

### Adaptor-dimer suppression using LNA antisense oligonucleotide

Library preparation protocol was identical as described above with the following modifications: L2 ligation and RT1 mixtures contained additional 10 μM antisense LNA oligo (final concentration). After washing and resuspending in mild wash buffer after reverse transcription reaction, the beads were incubated with 0.1 N NaOH (final concentration) and incubated at 95 C for 15 minutes. The reaction was neutralized with 0.1 volume of 1 M Tris (pH = 7.0).

### Preparing ribosomal footprints from mouse brains

Animal procedures were performed in compliance with national and international ethical guidelines and regulations, and were approved by the local animal welfare authorities at Heidelberg University Interfaculty Biomedical Research Facility (T-64/17). RjOrl:SWISS (RRID:MGI:5603077) mice (*Mus musculus*) were purchased from Janvier Labs (France). Mice were euthanized by cervical dislocation.

Frozen tissues were lysed in 150 μL of ice-cold lysis buffer (20 mM Tris-HCl, pH 7.5; 150 mM NaCl; 5 mM MgCl_2_; 1% (v/v) Triton X-100; 1 mM DTT; 0.4 U/mL Ribolock; and 100 μg/mL cycloheximide) using a micro pestle. Lysates were clarified by centrifugation at 20,000 × g for 7 min at 4°C. For nuclease digestion, 450 U RNase I (Ambion) and 3.75 U Turbo DNase I (Thermo Fisher Scientific) were added, and samples were incubated at 25°C for 45 min with gentle agitation. Digestion was stopped by the addition of 0.5 μL SUPERase In RNase Inhibitor (Ambion).

To purify ribosome-protected fragments, lysates were overlaid on a 0.7 mL 30% sucrose cushion in a 13 × 51 mm centrifuge tube (Beckman Coulter). Samples were centrifuged at 100,000 rpm for 1 hour at 4°C using an S100-AT6 rotor (Ultracentrifuge Sorvall Discovery M120 SE). The supernatant was discarded, and the pellet was resuspended in 700 μL of 10 mM Tris-HCl (pH 7.0). To extract RNA, 40 μL of 20% SDS and 750 μL of 65°C acid phenol-chloroform were added, followed by incubation at 65°C for 10 min with agitation. After centrifugation at maximum speed for 4 min, the aqueous phase was transferred to a fresh tube containing 700 μL acid phenol-chloroform, incubated at room temperature with intermittent vortexing and centrifuged for 4 min. Next, 600 μL chloroform was added, vortexed, and centrifuged for 4 min. RNA was precipitated overnight at −70°C in the presence of 600 μL isopropanol, 66.7 μL of 3 M sodium acetate (pH 5.5), and 2 μL GlycoBlue (Thermo Fisher Scientific). RNA was pelleted by centrifugation for 40 min at maximum speed, washed with 80% ethanol, and resuspended in 12.5 μL of 10 mM Tris-HCl (pH 7.0). Footprints were extracted and processed as described above with the following modifications: fragments ranging from 27 to 33 nt were excised and extracted with agitation at 70 °C for 10 minutes.

### rRNA depletion

We utilized a modified version of the previously published Ribocutter tool (26) to deplete rRNA from RP libraries. sgRNAs were designed to target the most abundant contaminants of a previously sequenced library derived from mouse telencephalon. To enhance the efficiency of rRNA removal, we used a lower library concentration (6 nM) as input for Cas9-mediated depletion and extended the Cas9 treatment to 4.5 hours. We incorporated an additional 6% PAGE-TBE step to remove preferentially amplified adapter-dimers following PCR re-amplification. All further steps were performed according to the original protocol.

### Preparation of samples for RNA-seq

CAG80099 cells (*E. coli* K-12 MG1655 *rpoC*::3xFLAG-kan, Carol Gross lab collection, UCSF) were grown overnight, diluted 1:100 in 100 ml of M9 medium supplemented with 0.4% glucose and 0.2% casamino acids and grown until OD_600_ ≈ 0.6. Cultures were poured onto 27.5 g of ice pre-chilled at −80 ºC and 10 ml of freshly made 30% formaldehyde were promptly added. Cells were crosslinked on ice with gentle shaking for 10 min and pelleted by centrifugation (15000 x g, 5 min, 4 ºC). The crosslinking reaction was quenched by resuspending the bacteria in 120 ml of ice-cold quenching buffer (50 mM HEPES pH 7.0, 100 mM NaCl, 10 mM MgCl_2_, 0.4% Triton X-100, 0.1% NP-40, 250 mM glycine) and gently shaking them on ice for 10 min. Cells were pelleted again (15000 x g, 5 min, 4 ºC) and resuspended in 45 ml of wash buffer (50 mM HEPES pH 7.0, 100 mM NaCl, 10 mM MgCl_2_, 0.4% Triton X-100, 0.1% NP-40). The pelleting and washing steps were repeated with 15 ml and 1 ml of ice-cold wash buffer. Bacteria were pelleted again (15000 x g, 5 min, 4 ºC) and resuspended in 725 μl of lysis buffer (50 mM HEPES pH 7.0, 100 mM NaCl, 10 mM MgCl_2_, 0.4% Triton X-100, 0.1% NP-40, 5 mM CaCl_2_, 100 U/ul DNase I (Roche, 04716728001), 1 mM PMSF). The cell suspension was flash-frozen in liquid nitrogen and bacteria were cryo-lysed in a mixer mill (Retsch MM 400) for 5 cycles of 3 min at 15 Hz with nitrogen-chilling steps between cycles. Cell powder was thawed at 37 ºC for 2 min and the lysate was nutated at 4 ºC for 15 min for DNA digestion. The lysate was cleared by centrifugation (4000 x g, 15 min, 4 ºC) and total RNA was quantified by Qubit™ (Thermo Scientific). 10 μg of RNA was aliquoted and diluted in wash buffer up to 695 μl in a safe-lock 1.5 ml tube. 15 μl of 0.5 M EDTA, 3 μl of 2.5 M glycine, and 37.5 μl of 20% SDS were added and the sample was mixed by inversion. 750 μl of pre-warmed acidic phenol-chloroform-isoamyl alcohol (Sigma, P1944) were added and cross-linking was reverted by incubation in a 65 ºC heating block for 45 min with 1500 rpm shaking. Samples were chilled on ice for 5 min and centrifuged (15000 x g, 5 min). About 700 μl of the aqueous phase were transferred to a fresh tube and 3 μl of GlycoBlue™ (Thermo Scientific, AM9515), 77.7 μl of 3 M NaOAc pH 5.5, and 780 ml of isopropanol were added. Samples were vortexed for about 15 s and incubated overnight at −20 ºC. RNA was pelleted by centrifugation (20000 x g, 1 h, 4 ºC) and the pellet was washed with 80% ice-cold ethanol. The pelleting and washing steps were repeated for up to three cycles. The RNA pellet was air-dried at 55 ºC for 3 min and resuspended in 16 μl of 10 mM Tris pH 7.0. RNA was quantified by Qubit™ (Thermo Scientific) and the RNA quality was tested by Bioanalyzer. rRNA was depleted with Pan-Prokaryote riboPOOL (siTOOLs Biotech) following the manufacturer’s guidelines and the sample was concentrated to 18 μl with RNA Clean & Concentrator Kit (Zymo Research). RNA was fragmented with NEBNext® Magnesium RNA Fragmentation Module (NEB) for 15 minutes and concentrated to 6 with RNA Clean & Concentrator Kit (Zymo Research). The fragmented RNA (2.4 μl) was directly used as an input for the library preparation protocol.

### Data processing and analysis

Demultiplex reads were processed as described earlier (22) with the following changes for libraries prepared with the new protocol:

The 3’ adaptor was trimmed with cutadapt (36) using the modified command

cutadapt command 1:

~~~
cutadapt -a AGATCGGAAGAGCACACGTCTGAACTCCAGTCAC -j 8 --nextseq-
trim=20 --discard-untrimmed -O 6 -m 25 -o output.fastq input.fastq.gz
~~~

After UMI extraction (8), 5’ adaptor was removed using cutadapt with the following command.

cutadapt command 2:

~~~
cutadapt -g ^CCCCCC -g ^CCCCC -g ^CCCC -m 10 -M 70 -o output.fastq.gz
input.fastq.gz
~~~

To remove contaminating sequences, trimmed reads were mapped to the indexed libraries of *E. coli-*specific, yeast-specific rRNA (provided in the supplementary material) and aligned with bowtie (37) 1.3.1 using the following command

~~~
bowtie --threads 8 -t -n 2 –best input.fastq.gz --un rRNA_depleted.fastq
~~~

or Mouse-specific contaminating RNAs were obtained from RNAcentral (rRNAs, mtRNAs, tRNAs) using Bowtie2 v.2.5.1 (parameters: --phred33 -L 20 -N 1 -t --no-unal).

The trimmed reads were aligned to *Escherichia coli* genome (build GCA_000499485.1), *Saccharomyces cerevisiae* genome (build GCA_000146045.2) or *Mus musculus* (GRCm39, GCA_000001635.9)) with bowtie 1.3.1 (*E. coli* and yeast) and STAR aligner (38) v.2.7.11a (mouse). Further processing was done with custom Python and R scripts available in the supplementary materials.

---

**Figure S1:**
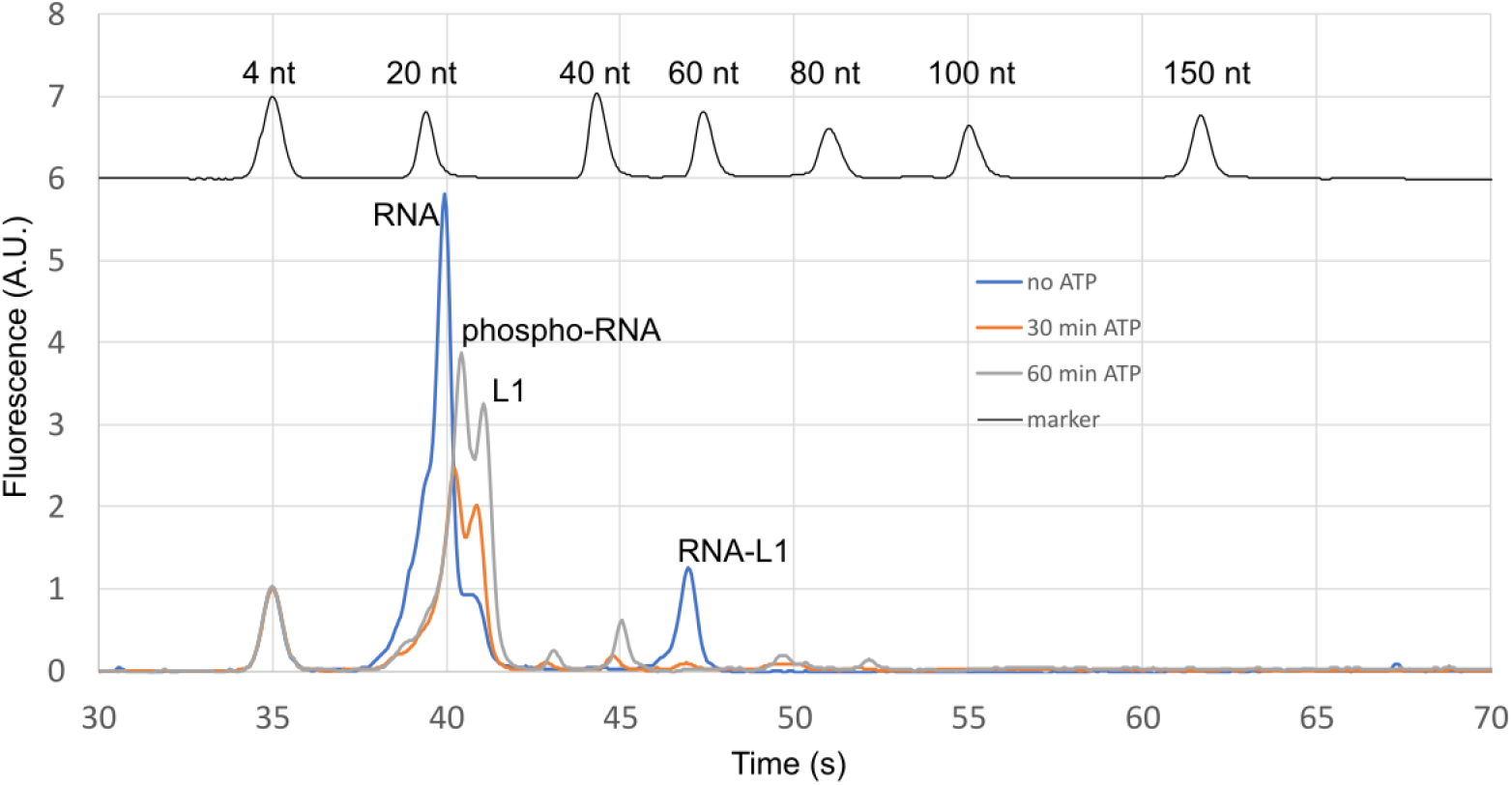
Bioanalyzer traces of L1 reaction products based on different phosphorylation conditions of RNA. Phosphorylation/dephosphorylation was carried out (i) without ATP for 60 minutes (no ATP) or (ii) without ATP for 30 minutes after which 1 mM ATP was added for additional 30 minutes (30 min ATP) or (iii) with 1 mM ATP for 60 minutes (60 min ATP). L1 ligation was carried out for 1 hour, after which the products were precipitated with isopropanol in the presence of NaOAc and Glycoblue, resuspended in 10 mM Tris (pH = 7.0) and loaded on Bioanalyzer small RNA chip. Traces were normalized to small RNA marker (4 nt). RNA-L1 peak corresponds to the expected ligation product.

**Figure S2:**
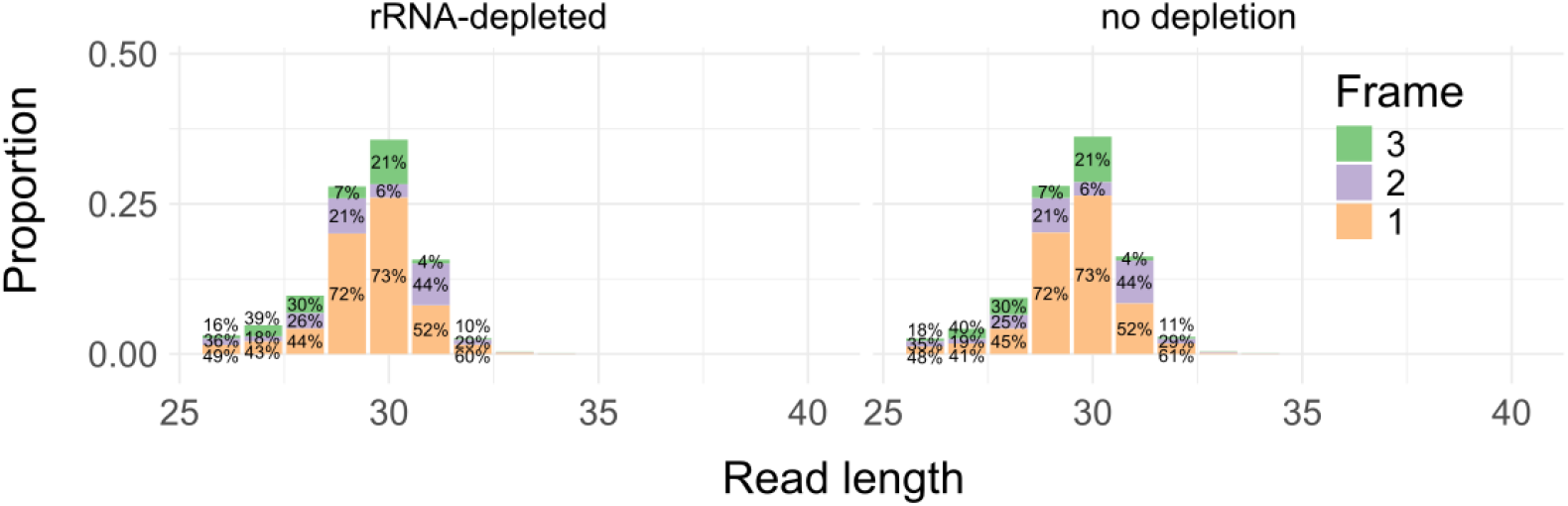
Cas9-mediated depletion has no impact on footprint length distribution and detected periodicity of cds-aligned reads in mouse brain samples. For each footprint length, footprints assigned to each frame are listed in ascending order, i.e. Frame 3 on top and Frame 1 on the bottom.

